# Bridging the gap between genome-wide association studies and network medicine with GNExT

**DOI:** 10.64898/2026.01.30.702559

**Authors:** Lis Arend, Fabian Woller, Bastienne Rehor, David Emmert, Johannes Frasnelli, Christian Fuchsberger, David B. Blumenthal, Markus List

## Abstract

**Motivation:** A growing volume of large-scale genome-wide association study (GWAS) datasets offers unprecedented power to uncover the genetic determinants of complex traits, but existing web-based platforms for GWAS data exploration provide limited support for interpreting these findings within broader biological systems. Systems medicine is particularly well-suited to fill this gap, as its network-oriented view of molecular interactions enables the integration of genetic signals into coherent network modules, thereby opening opportunities for disease mechanism mining and drug repurposing.

**Results:** We introduce GNExT (GWAS Network Exploration Tool), a web-based platform that significantly extends the scope of exploration of variant-level effects and significance beyond those provided by existing solutions. By including MAGMA and Drugst.One, GNExT allows its users to study genetic variants in the context of the latest systems medicine approaches, extending to the identification of potential drug repurposing candidates. Moreover, GNExT advances platform implementation well beyond the current state of the art by offering a highly standardized Nextflow pipeline for data import and preprocessing, allowing researchers to deploy their study results on a sophisticated web interface with minimal implementation overhead. We demonstrate the utility of GNExT using a genome-wide association meta-analysis of human olfactory identification, in which the framework translated isolated GWAS signals to potential pharmacological targets in human olfaction. Furthermore, the deployment of a GNExT instance on European-ancestry Pan-UK Biobank data demonstrates the framework’s scalability, resulting in a comprehensive large-scale resource encompassing thousands of traits and enabling new network medicine-based investigations.

**Availability and Implementation:** The complete GNExT ecosystem, including the Nextflow preprocessing pipeline, the backend service, and frontend interface, is publicly available on GitHub (https://github.com/dyhealthnet/gnext_nf_pipeline, https://github.com/dyhealthnet/gnext_platform). The public instances of the GNExT platform on olfaction and Pan-UKBB are available under https://olfaction.gnext.gm.eurac.edu and https://panukbb-eur.gnext.gm.eurac.edu.

## 1. Introduction

Advances in genotyping, together with widespread electronic health records and large population-based cohorts, have transformed the scale of genetic studies [37]. Modern datasets now comprise genetic profiles spanning millions of loci for millions of individuals across thousands of traits, enabling genome-wide association studies (GWAS) of unprecedented breadth [65]. These high-dimensional GWAS resources underpin diverse downstream analyses and have substantially advanced our understanding of the genetic architecture and biological mechanisms of human traits and diseases [65, 67, 58].

Despite the growth of GWAS data, standard single variant association tests fail to fully capture the genetic architecture of complex traits, which are highly polygenic and driven by numerous variants of small individual effects [60, 17, 68, 52]. To control genome-wide false positives under this high-dimensional testing burden, very stringent significance thresholds are required, which severely reduce power to detect weak but genuine associations unless sample sizes reach several hundred thousand individuals [68, 26, 52]. To address these limitations and capture the biological architecture underlying complex traits, gene-based approaches have gained prominence [49, 32]. By aggregating numerous weak variant-level signals within genes or genomic regions into composite gene-level statistics, these methods reduce the multiple testing burden, capture combined genetic effects, and increase power to detect genuine disease contributors [49, 17]. While methods such as MAGMA [17], VEGAS [38], and PascalX [33], enhance statistical power, they primarily identify associations without fully resolving the biological context underlying these signals. Bridging this gap requires combining association results with mechanistic insights, a task for which systems medicine and network-based approaches provide essential frameworks [25].

In systems medicine, increasing attention is being directed toward the observation that disease-associated genes are not randomly distributed in biological networks but cluster within specific regions of the human interactome. These genes, typically termed seed genes, and the interactions of their encoded proteins with neighboring molecules are used to expand the initial seed set and define disease modules that represent the underlying pathogenic mechanisms [46, 25]. Beyond providing mechanistic insight, disease modules form the basis for network pharmacology strategies that systematically identify therapeutic targets by considering both seed and neighboring nodes [50]. This enables the discovery of targets that can be modulated by existing drugs and thus supports drug repurposing, an attractive alternative to de novo drug development given its reduced cost and shorter timelines [54, 34, 29].

In parallel, the landscape of post-GWAS analyses has rapidly evolved to accommodate the increasing complexity and diversity of genetic research needs [32]. Broadly, existing resources fall into three categories. The first category consists of functional annotation and interpretation tools, including platforms such as FUMA [70], postGWAS [69], GWAShug [12], and CTG-VL [16]. These tools allow users to upload, explore, and interpret summary statistics on a per-trait basis. FUMA, specifically, is a web-based interface enabling the comprehensive functional mapping and annotation of GWAS results by integrating multiple biological databases and supporting users in annotating, prioritizing, visualizing, and interpreting associations for individual phenotypes. Notably, platforms such as FUMA and CTG-VL also offer gene-based association testing with MAGMA and fastBAT [5], respectively. Several of these platforms also host curated analyses in which post-GWAS results are made publicly accessible. Nevertheless, they are designed for post-GWAS analyses on a single trait at a time, reflecting their emphasis on detailed, phenotype-specific investigation. The second category encompasses global or study-specific software tools primarily designed for GWAS data dissemination. While the GWAS Catalog [58] and openGWAS [20] serve as central repositories for curated GWAS results, the GWAS Atlas [39] extends functionality by enabling interactive exploration of curated GWAS and post-GWAS results. Study-specific portals, including the GWAS Explorer [41] and the collection of Human Genetics Amplifier (HuGeAMP) knowledge portals, such as the Cardiovascular Disease Knowledge Portal [15], are designed to focus on specific cohorts or disease areas. The third category consists of visualization frameworks like PheWeb [24] and its successor PheWeb 2 [7], which enable the exploration of large-scale, population-specific GWAS datasets through interactive visualization, navigation, and publication of GWAS and PheWAS results. A distinct feature of PheWeb is its support for deploying independent instances, allowing researchers to operate the platform on their own datasets without the need to redesign study-specific frameworks. This capability has already supported several large-scale Biobank and population genetics resources [10, 35, 73, 47]. While PheWeb 2 extends the original PheWeb by functionalities enabling exploration and comparison of stratified GWAS summary-level data, further downstream post-GWAS analyses, such as gene-based tests, are not integrated.

Despite the breadth of available tools operating on GWAS summary statistics, most platforms remain centered on trait-specific visualization and exploration, offering limited integration of systems-level concepts. As a result, these tools provide limited insight into how genetic associations converge into coherent biological processes, and do not directly support the translation of these mechanistic insights into therapeutic hypothesis generation, an area where network medicine offers substantial advantages.

Building on principles of interactive, scalable, and decentralized deployment, we introduce GNExT (GWAS Network Exploration Tool), an integrated framework that in addition to providing first order visualization and interpretation of GWAS results via PheWeb, LocusZoom, and MAGMA, extends this paradigm by integrating GWAS summary statistics with network medicine methodologies (Figure 1). A dedicated, configurable Nextflow-based preprocessing pipeline, which processes GWAS data together with user-defined configuration parameters, substantially streamlines and automates data preparation, making deployment accessible to researchers who may lack computational expertise, while also enabling data extensions beyond existing deployments. Variant-level association statistics are aggregated into gene-level metrics using MAGMA, and all generated output files are consolidated into a directory that serves as the primary input for deploying the GNExT platform. The GNExT platform can be deployed via a docker-compose stack and provides variant-, gene-, and trait-centric visualization for exploratory analyses. Potential risk genes can be identified by applying significance thresholds to gene-level *P*-values. These genes serve as seed nodes for detecting disease modules in protein-protein interaction networks through the Drugst.One interface [42]. GNExT thus facilitates the elucidation of candidate disease mechanisms and the prioritization of potential therapeutic targets directly from GWAS summary statistics. We demonstrate the use of GNExT through two use cases, one focusing on an exemplary olfaction-focused GWAS study, while the other one shows how GNExT can scale to larger scale in PanUK Biobank. Together these use cases show the versatile applications of GNExT.

**Figure 1.**
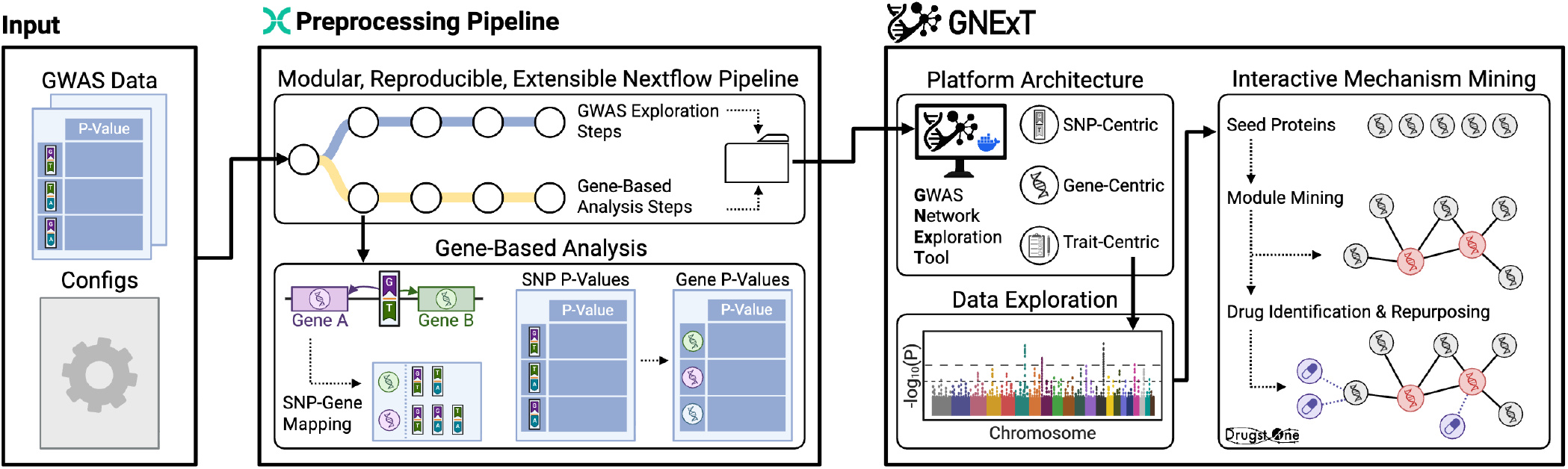
Overview of the GNExT framework. Input data consists of GWAS summary statistics together with study-specific configuration parameters. A Nextflow-driven preprocessing pipeline transforms and organizes the data in a scalable way, ultimately generating the primary input directory required for platform deployment. GNExT extends its functionalities beyond data exploration by linking GWAS results to network medicine through gene-based analyses and an integrated, interactive mechanism mining interface. Created with BioRender.com.

## 2. Results

### 2.1. Nextflow preprocessing pipeline allows for efficient and flexible data preparation

Our Nextflow-based pre-processing pipeline [19] provides a modular and extensible workflow architecture that orchestrates and parallelizes workflow execution and allows resuming computations after an error without repeating previously completed, computationally intensive steps. Likewise, it supports incremental dataset expansion by enabling the addition of new traits while reusing previously generated trait-specific results, avoiding redundant computation. The pipeline is widely usable as it supports local execution through Conda, containerized environments via Docker or Singularity, and distributed execution using SLURM-based HPC systems, ensuring flexibility, scalability, and reproducibility for large-scale genomic analyses.

The Nextflow pipeline prepares GWAS summary data for integration into the GNExT platform, including data harmonization, variant annotation using the Ensembl Variant Effect Predictor (VEP) ([45]), and data processing into Lightning Memory-Mapped Database (LMDB) structures to enable efficient downstream querying (Figure 2). In addition, analogous to PheWeb, the pipeline generates preprocessed inputs for standard visualizations and tabular outputs, including Manhattan, Q-Q, and PheWAS plots as well as a Top Hits table. A critical step in the pipeline is the use of MAGMA to perform gene-based association testing. By aggregating SNP-level summary statistics while accounting for local linkage disequilibrium (LD), MAGMA generates gene-level p-values that allow us to statistically prioritize candidate risk genes. These prioritized genes then serve as the foundational seeds for downstream network medicine analysis, such as module discovery and drug repurposing (see Methods for details).

**Figure 2.**
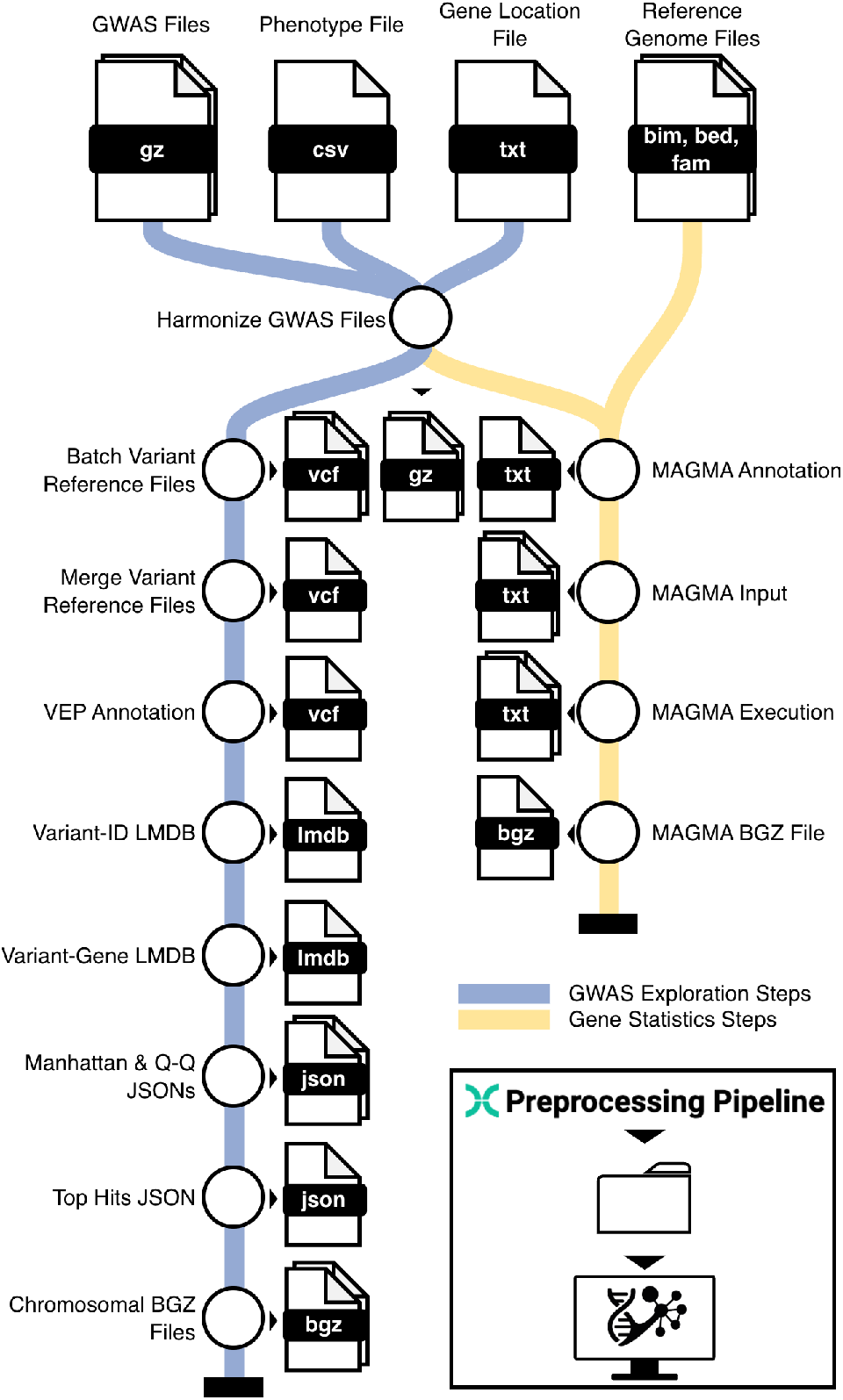
Overview of the Nextflow-based preprocessing pipeline. The pipeline comprises two main stages. First, GWAS data are harmonized, and a reference VCF containing all unique variants is assembled for downstream annotation. Two LMDB databases are generated to store variant-rsID and variant-gene mappings. The workflow also produces JSON files for Manhattan and Q-Q plots, a Top Hits table, and per-chromosome data files for efficient variant-level retrieval. If enabled by user configuration, MAGMA input data are prepared, and gene-level association statistics are computed. All outputs generated by the preprocessing pipeline are consolidated into a deployment directory, which forms the essential data backbone for launching a GNExT instance.

### 2.2. GNExT application connects GWAS with network medicine in a decentralized manner

The GNExT backend and frontend receive the output of the Nextflow pre-processing pipeline along with a set of configuration parameters (see Methods section).

With GNExT, users can query variant-, trait-, or gene-level result pages, analogous to PheWeb (Supplementary Figure 1A). In addition, a dedicated Top Hits page summarizes the most significant variants identified across all traits represented within the platform, offering a concise overview of key associations. The variant-specific page provides detailed Ensembl VEP annotations, allele frequency comparisons with 1000 Genomes [1] and gnomAD [13], cross-references to external databases, and nearest-gene assignments (Supplementary Figure 1B). An integrated LocusZoom-based PheWAS view [9], accompanied by a summary table, presents all trait associations for the variant with interactive filtering options. Within the interface, selecting a gene symbol redirects to the gene-specific page, which lists the most significant proximal variant for each trait and provides a trait-specific LocusZoom plot displaying regional association signals, LD patterns, and gene annotations (Supplementary Figure 1C).

The trait-specific page visualizes all association signals from a single GWAS through interactive Manhattan and Q–Q plots and includes an extended Top Hits table with newly introduced retrieval options that filter results by *P*- value, genomic window, or chromosomal region. Gene-level associations from MAGMA are displayed analogously, and significant genes (default Bonferroni-corrected *P*-value < 0.05) can be selected as seeds and transferred to the network medicine interface for downstream analyses (Supplementary Figure 2A).

The network medicine-specific page allows users to select a predefined seed gene list and transfer the seed genes to the integrated Drugst.One interface (Supplementary Figure 2B). Drugst.One provides tools for disease module refinement, drug repurposing, and additional analyses such as functional enrichment (see Methods). Users may choose among multiple biological network sources, including seven protein-protein, five drug-protein, three protein-disease, and three drug-disease interaction databases, with NeDReX ([56]) selected by default as this knowledge graph integrates more than twenty different data sources on drug, gene, protein, and other interactions. Users can also navigate to the gene-specific pages to inspect GWAS signals for seed genes or newly identified module genes.

### 2.3. Linking olfactory GWAS signals to systems-level mechanisms with GNExT

To demonstrate how GNExT bridges the gap between statistical associations and mechanistic understanding, we applied the framework to a recent GWAS meta-analysis of human olfactory identification [23]. The GNExT platform on olfaction data, accessible at https://olfaction.gnext.gm.eurac.edu, comprises 39 traits, 23,544,328 unique genetic variants, and 18,547 genes. While GWAS successfully identifies genetic loci associated with traits, interpreting these findings in the context of biological systems remains a challenge. Here, we utilized GNExT to move beyond isolated variant lists to reconstruction of signaling pathways and potential drug interactions underlying olfactory function.

We utilized summary statistics from the “score all” phenotype (*n* = 21,495), a global measure of olfactory ability derived from the Sniffin’ Sticks identification test. Initial exploration via GNExT confirmed the strong signal on chromosome 11 (designated as locus 8 in [23]) (Figure 3A). To resolve these variant-level signals into interpretable biological units, we employed MAGMA within the Nextflow preprocessing pipeline, identifying 33 significant seed genes (Bonferroni-corrected *P*-value < 0.05), predominantly in olfactory receptor (OR) gene families OR5 (17 genes), OR8 (11 genes), and OR9 (2 genes), with additional associations in OR10AG1, OR9G4, and TRIM51 (Supplementary Figure 2A and Supplementary Table 1). While identification of these candidate genes offers some potential insights into the genetic architecture of olfaction, examining receptors in isolation offers limited insight into the downstream signaling machinery that translates chemical binding into neuronal perception.

**Figure 3.**
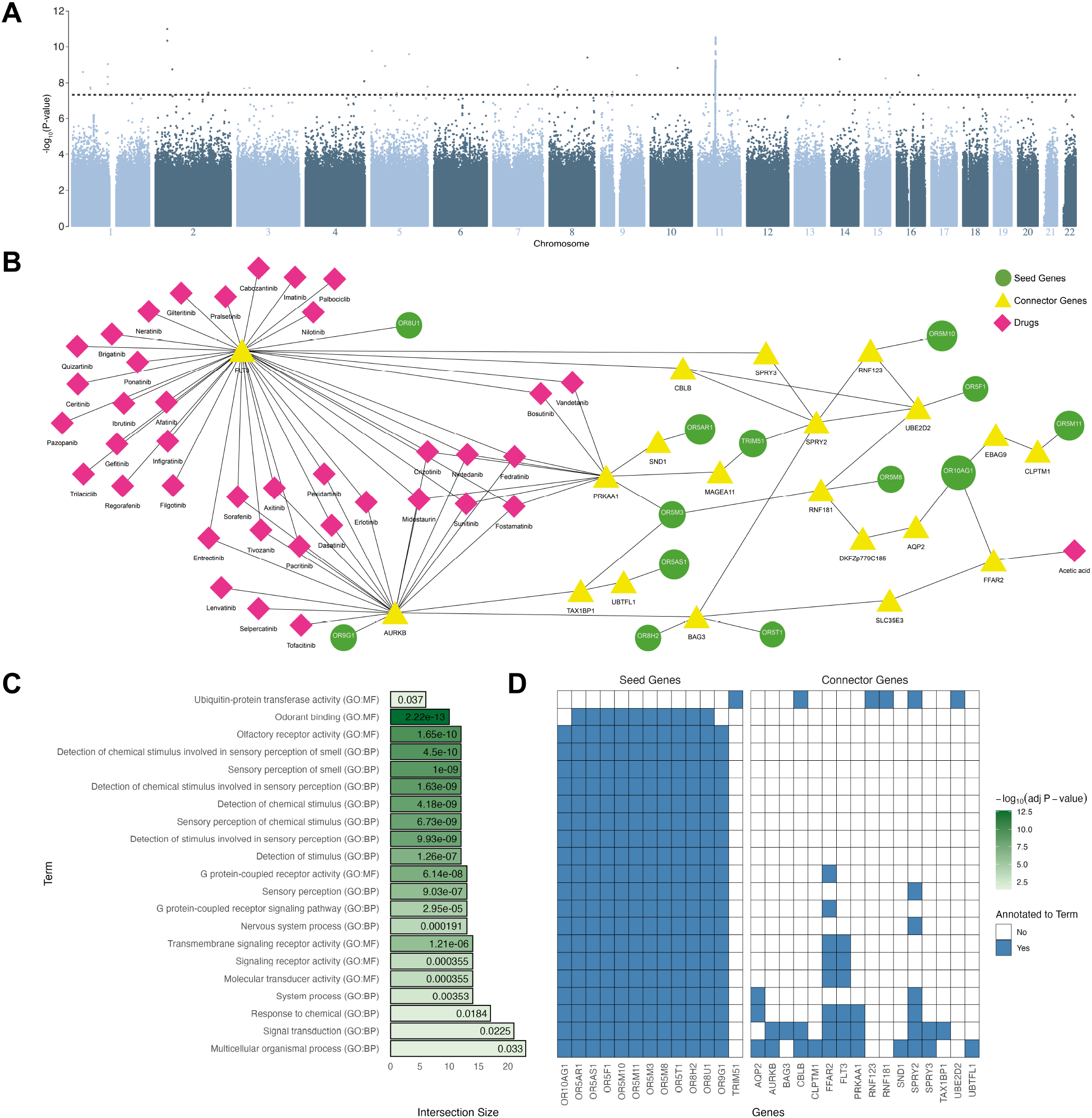
Results of the olfaction use case. (A) Manhattan plot from the GNExT olfaction instance, revealing a strong genome-wide significant association signal on chromosome 11. (B) Network module inferred using a Multi-Steiner Tree algorithm, consisting of MAGMA-identified significant genes (seed genes) and interacting genes required to connect the seed genes (connector genes). Drug prioritization was performed using harmonic centrality. (C) Functional enrichment analysis of the network module genes using g.Profiler, with significantly enriched terms ranked by intersection size, bars colored according to the −*log*_10_ adjusted *P*-value, and the corresponding adjusted *P*-values displayed within the bars. (D) Mapping of seed and connector genes to the enriched functional terms identified in (C). Four connector genes, namely MAGEA11, SLC35E3, EBAG9, DKFZp779C185, did not contribute to the statistical significance of any enriched terms; consequently, only 13 seed and 16 connector genes are shown.

To reconstruct the functional context of these receptors, we transferred these 33 (seed) genes into the Drugst.One network medicine environment within GNExT. Three of the 33 genes were not present in the underlying NeDReX database [56] and therefore could not be incorporated (Supplementary Figure 2B and Supplementary Table 1). We hypothesized that while ORs drive the genetic signal, they must physically interact with downstream signal transducers to effect perception. Using the “Quick drug search” option, triggering the sequential execution of the Multi-Steiner tree module detection and closeness centrality algorithm for drug prioritization, we identified a coherent network module comprising 13 out of the 30 integrated seed genes and 20 additional connector genes, i. e., proteins not significant in the GWAS itself but essential for maintaining network integrity (Figure 3B). Functional enrichment of the network module with g.Profiler [55] highlighted the significant enrichment in molecular function (MF) and biological processes (BP) terms related to olfaction, including “Odorant binding”, “Olfactor receptor activity”, and “Detection of chemical stimulus involved in sensory perception of smell” (Figure 3C). Consistent with expectations, all seed genes except TRIM51 contributed to the olfaction-associated enriched terms (Figure 3D). In turn, connector genes such as FLT3, AURKB, PRKAA1, or FFAR2 contributed to terms associated with olfactory transduction downstream from odorant binding, such as “G-Protein coupled receptor signaling pathways”, “molecular transducer activity”or “signal transduction”. This aligns with the known biology of olfactory perception, where ORs (GPCRs) rely on downstream cascades involving adenylate cyclases and kinase activity to transmit signals to the olfactory bulb.

Finally, we asked whether this systems-level view could generate actionable pharmacological insights. By querying the expanded network for drug-gene interactions using harmonic centrality, we identified 39 drugs targeting the connector genes, specifically FLT3, AURKB, PRKAA1, or FFAR2 (Figure 3). In our network, FLT2, AURKB, and PRKAA1 were connected to OR8U1, OR9G1, and OR5M3, respectively. Of these, only OR5M3 has been deorphanized and responds to furaneol, which gives different foods a caramel-note [28]. The drugs connected to FLT2, AURKB, and PRKAA1 belong to the group of tyrosine kinase inhibitors (TKI), which target tyrosine kinases that govern cell growth and division and are used in the treatment of various types of cancer [62, 72, 51]. Crucially, this network-derived finding offers a mechanistic hypothesis for a known clinical phenomenon: TKIs are associated with olfactory dysfunction such as anosmia, hyposmia, and parosmia [18, 22, 66, 6]. Furthermore, tyrosine kinase levels in the olfactory bulb play a role in odor memory consolidation, which may be a potential pathway by which these drugs affect olfactory perception [64]. Through this workflow, GNExT successfully translated raw GWAS summary statistics into a plausible mechanistic hypothesis, linking genetic variants to a specific class of drugs known to perturb the phenotype.

### 2.4. GNExT scales to thousands of traits

To further assess the scalability of the platform, we deployed a GNExT instance on Pan-UK Biobank data of European ancestry comprising 7,160 traits. This large-scale setting, characterized by a high-dimensional spectrum of traits and associated GWAS summary statistics (2.1 TB), served as a stress test for both the preprocessing pipeline and the interactive exploration components of GNExT. To demonstrate that the ecosystem can robustly accommodate cohort-scale resources beyond small panel analyses, we used Nextflow’s built-in functionality to track and report runtime and memory metrics for the individual processes. The reported benchmarks for both the olfaction and Pan-UKBB use cases exclude the time required to download variant annotation cache data, as these resources were already available locally, as well as the time and memory needed to create the required conda environments.

For the olfaction use case, Ensembl VEP annotation represented the most time-consuming preprocessing step in terms of CPU usage (Supplementary Figure 3). Additional stages, including GWAS harmonization, Manhattan and Q-Q JSON file generation, chromosome-specific BGZ file construction, and MAGMA analyses, each surpassed one hour of accumulated CPU time. Nevertheless, the overall wall-clock runtime of the pipeline was markedly reduced through the systematic use of batching and parallelization of 69 jobs in total. The complete workflow finished in approximately 11 hours, leveraging SLURM-based parallelization and the configuration parameters provided in the corresponding GitHub repository.

For the Pan-UKBB platform, 273 SLURM jobs were executed in total, and the pipeline operated for around 7 days to complete data preprocessing under the specified configuration parameters.

The trait-specific steps, with traits organized into 48 batches, including GWAS data harmonization, batch variant reference file creation, the Manhattan and Q-Q JSON file generation, as well as MAGMA input and test execution, accounted for a substantial proportion of the overall runtime, analogous to the olfaction runtime but scaled to the markedly larger number of phenotypes (Figure 4). The chromosomal BGZ file creation was particularly time-consuming, as this step required querying 7,160 GWAS summary statistics files, rather than 39, to construct a variant-centric representation of the results. In addition, we observed pronounced variability in MAGMA execution times across trait batches and in chromosomal BGZ file creation times, the latter attributable to differences in chromosome sizes.

**Figure 4.**
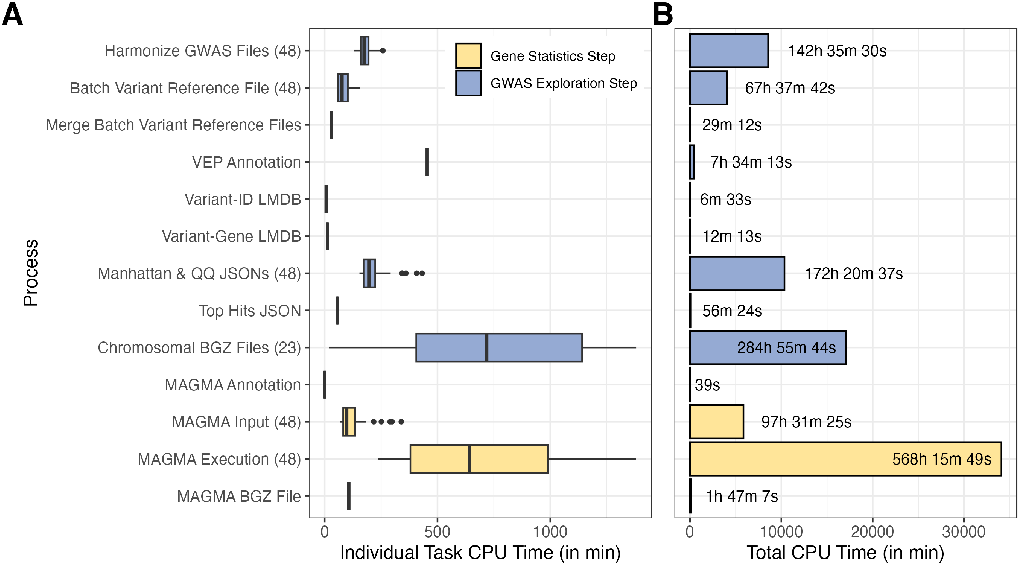
Individual and total CPU time per process of the Nextflow preprocessing pipeline for the Pan-UKBB use case. Individual task CPU time was recorded intrinsically by Nextflow (A) and aggregated across all jobs belonging to each process (B). The x-axis denotes the process types, with the corresponding number of individual jobs per process arising from trait- or chromosome-batching shown in brackets.

Finally, for the Nextflow preprocessing pipelines for both use-cases we assessed the peak resident set size (RSS), representing the maximum physical RAM occupied during execution. Unexpectedly, the mean peak RSS varied substantially across both pipeline steps and use cases (Supplementary Figure 4). For the olfaction data, MAGMA input generation showed the highest memory consumption, followed by the harmonization of GWAS files. Memory usage during MAGMA input preparation even exceeded that observed in the Pan-UKBB use case, as the reference variant set used for MAGMA contained a larger number of variants that were fully loaded into RAM at this stage to intersect with the GWAS variants. In the Pan-UKBB use case, chromosomal BGZ file generation, the most computationally intensive step (Figure 4), demonstrated substantially higher memory consumption than all other processes. This elevated memory requirement stems from simultaneously loading 7,160 variant-level statistics into memory, compared to only 39 phenotype-specific statistics in the olfaction dataset. Other pipeline steps, including data harmonization, variant reference file generation, and VEP annotation, exhibited comparable memory usage across both use cases despite the difference in dataset scale.

To summarize, these results demonstrate that the GNExT ecosystem is applicable across diverse dataset scales, from small-scale phenotype studies to large phenotype repositories spanning thousands of traits. While large-scale applications require substantial computational resources (e.g., HPC clusters with sufficient memory and CPU allocations), such infrastructure is typically available in research environments that routinely generate, maintain, and analyze datasets of this magnitude.

### 2.5. GNExT on Pan-UKBB European ancestry data represents a large-scale resource for GWAS interpretation

The Pan-UKBB European ancestry GNExT instance, accessible at https://panukbb-eur.gnext.gm.eurac.edu, constitutes a robust resource for future exploratory analyses as it integrates 7,160 traits, comprising 23,118,158 unique genetic variants, mapping to 20,104 genes (Figure 5A). Following the Pan-UKBB trait taxonomy, these traits are grouped into 6 categories: biomarkers, categorical, continuous, icd10, phecode, and prescriptions. Categorical traits comprise the largest group, whereas biomarkers represent the smallest (Figure 5B). GWAS analyses were conducted in more than 400,000 European-ancestry samples across over 4,500 traits, underscoring the large-scale nature of this resource in both phenotypic breadth and cohort size (Figure 5C).

**Figure 5.**
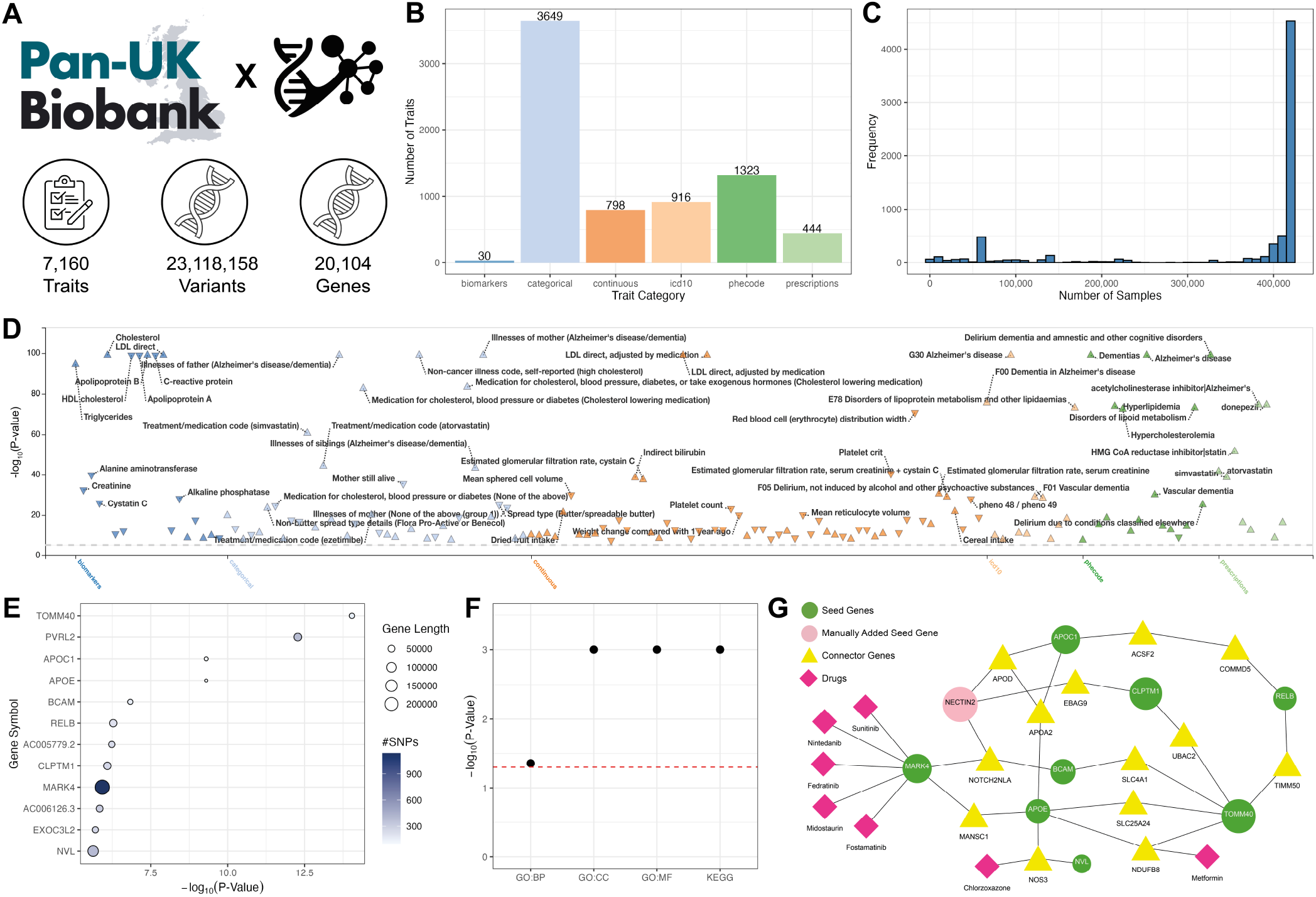
Results of the European-ancestry Pan-UKBB use case. (A) Overview of the integrated resource, summarizing the number of traits, unique genetic variants, and mapped genes. (B) Distribution of traits across Pan-UKBB-defined phenotypic categories. (C) Distribution of sample sizes employed across all included GWAS analyses. (D) PheWAS plot generated within GNExT for variant rs429358 (chromosome 19; position 45,411,941; T>C), filtered at the genome-wide significant threshold of *P*-value < 5 × 10^−8^. (E) Gene-level association results derived from MAGMA for the Alzheimer’s disease phecode (trait ID: phecode-290.11-both sexes), displaying statistically significant genes (Bonferroni-corrected *P*-value < 0.05). (F) DIGEST-based functional coherence evaluation results of the network module inferred via the Multi-Steiner tree module detection algorithm using the “Quick drug search” functionality. (G) Network module including MAGMA-identified significant genes (seed genes) and interacting genes required to connect the seed genes (connector genes), with drug prioritization performed using harmonic centrality.

Alzheimer’s disease, the most prevalent form of dementia, serves as a compelling case study within this framework, given its well-characterized genetic architecture, spanning common genetic variants alongside rarer pathogenic mutations. Among genetic loci implicated in Alzheimer’s disease risk, the Apolipoprotein E (APOE) gene, the predominant cholesterol transport protein in the central nervous system, was among the earliest to be identified [61, 14]. Specifically, the single nucleotide variant rs429358 (chromosome 19; position 45,411,941; T>C), a well-characterized variant within the APOE locus, represents one of the strongest common genetic risk factor for late-onset Alzheimer’s disease [8, 30]. Leveraging GNExT, we investigated this variant in greater detail to characterize the breadth of phenotypic associations linked to this locus (Figure 5D). The phenotypic association landscape reveals robust signals across different dementia-related traits and lipid metabolism phenotypes, such as low-density lipoprotein (LDL) cholesterol levels, consistent with APOE’s established role in lipid metabolism [71].

Beyond variant-centric analyses, GNExT also enables trait-centric investigations, thus supporting both in-depth exploration of the genetic architecture underlying a specific phenotype and its potential for drug repurposing. Analogous to the approach employed in the Olfaction use case, we applied MAGMA to aggregate the variant-level association signals into gene-level statistics for the Alzheimer’s disease phecode (trait ID: phecode-290.11-both sexes), identifying 12 genes surpassing the significance threshold (Bonferroni-corrected *P*-value < 0.05) (Figure 5E). The mapping of MAGMA-derived gene-level signals onto the NeDReX network via the Drugst.One interface yielded 9 directly mappable genes, with PVRL2 requiring manual mapping via the “Edit Network” functionality of Drugst.One, as its alias NECTIN2. Leveraging the integrated “Quick Drug Search” functionality, the resulting disease module comprised 9 seed genes (EX0C3L2 could not be connected) and 13 connector genes. To assess the biological validity of the candidate disease module, we employed the DIGEST linkout within Drugst.One, which evaluates functional coherence through pairwise comparisons of gene set annotations from the Gene Ontology and the Kyoto Encyclopedia of Genes and Genomes (KEGG). While significant *P*-values were obtained across all three types of GO term categories, as well as KEGG pathways, the functional coherence signal was least pronounced for GO:BP (Figure 5F).

Finally, we analyzed drug-protein interactions to assess whether this systems-level perspective could yield actionable therapeutic insights into Alzheimer’s disease. Applying harmonic centrality-based drug prioritization, we identified seven candidate drugs, of which five target the same seed gene, namely MARK4, while two target connector genes within the disease module (Figure 5G). Metformin and Chlorzoxazone ranked highest among all candidates, targeting the connector genes NDUFB8 and NOS3, respectively. Metformin is a first-line pharmacological treatment for type 2 diabetes mellitus, primarily acting through AMPK-mediated inhibition of hepatic gluconeogenesis to reduce fasting blood glucose levels [36]. Notably, clinical evidence has suggested that Metformin use is associated with a reduced risk of developing Alzheimer’s disease and improved cognitive performance [11, 36, 44], positioning it as a compelling repurposing candidate. Chlorzoxazone is an FDA-approved muscle relaxant primarily used to treat musculoskeletal pain and stiffness. Beyond its classical indication, preclinical evidence has demonstrated its potential relevance to Alzheimer’s disease [4, 43]. The five drugs, Sunitinib, Nintedanib, Midostaurin, Fostamantinib, and Fedratinib are all multi-target tyrosine kinase inhibitors, primarily approved for oncological indications, whose identification through the MARK4 seed gene is mechanistically coherent, as MARK4 overexpression causes early phosphorylation of tau at Ser262, driving the formation of neurofibrillary tangles, a core hallmark of Alzheimer’s disease [48, 40, 63]. A multi-omics study aiming to study the possibility of repurposing approved anticancer drugs for Alzheimer’s disease treatment proposed Nintedanib and Sunitinib as potential repurposing drugs for Alzheimer’s disease [3].

Collectively, these findings demonstrate that the GNExT ecosystem can surface biologically plausible and clinically relevant drug repurposing hypotheses directly from GWAS-derived gene-level signals, underscoring the value of network-based, systems-level approaches in bridging genomic discovery and actionable therapeutic insights. Beyond this Alzheimer’s disease use case, the Pan-UKBB instance constitutes a comprehensive and openly accessible resource spanning thousands of traits and millions of genetic variants, inviting the broader research community to conduct analogous variant- or trait-centric analyses across the full phenotypic landscape captured by the GNExT instance on Pan-UKBB data.

## 3. Discussion

The GNExT ecosystem provides researchers with a streamlined framework for deploying a web-based platform that enables comprehensive exploration and visualization of GWAS summary statistics, while incorporating network medicine methodologies.

Specifically, through the application of MAGMA and Drugst.One, GNExT is the first web-based platform that seamlessly connects GWAS data and network medicine. To support both scientific analyses and dissemination, GNExT supports the generation of customizable tabular reports, summary statistics, and publication ready visualizations, thereby enabling researchers to efficiently interrogate results and export figures and data tables suitable for manuscripts and presentations.

To further ensure an interactive user interface while minimizing deployment effort and broadening accessibility, we developed a Nextflow-based pipeline for data preparation and preprocessing. Through Nextflow’s built-in workflow orchestration and caching mechanisms, all pipeline components are executed efficiently and reproducibly, enabling both streamlined deployment and efficient extension of existing platforms.

By applying GNExT to a GWAS meta-analysis of human olfactory identification and to an Alzheimer’s use case in European-ancestry Pan-UKBB data, we demonstrate that the framework facilitates the interpretation of variant-level association signals by contextualizing them into signaling pathways and prioritizing putative drug-target interactions relevant to the phenotype of interest. In addition, the application of the GNExT ecosystem to the large-scale Pan-UKBB dataset, encompassing over 7,000 traits, demonstrates its scalability to cohort-level resources, provided adequate computational infrastructure is available.

A limitation in the current GNExT ecosystem implementation has to do with the use of MAGMA for execution of gene-based testing. The constrained genomic windows employed with this method fail to capture long-range regulatory interactions, which can span hundreds of kilobases and are increasingly recognized as critical contributors to gene regulation [27]. In contrast, expanding flanking regions introduces substantial noise by assigning variants to unrelated genes, thereby diluting true biological signals. Integrating experimentally derived regulatory interaction datasets, such as eQTL or Hi-C datasets, can mitigate these issues [27]. Consequently, enabling more flexible, user-defined variant-to-gene mapping strategies represents an important avenue for enhancing future versions of GNExT.

Additionally, the Pan-UKBB results suggest several targeted optimizations of memory- and runtime-intensive processes within the Nextflow pipeline. Analogous to optimization opportunities identified in the MAGMA input generation step, the chromosomal BGZ file generation process could be refactored to accommodate substantially larger repositories (> 10.000 GWAS files) by adopting memory efficient data structures and introducing parallelization beyond chromosome level batching. Further, instead of re-annotating the full variant set across runs that differ in the included phenotypes, the pipeline could identify the subset of newly introduced variants by comparing the unique variant set between runs, annotating only these additions. Similarly, existing chromosomal BGZ files could be extended with new trait data rather than regenerated entirely. While many pipeline steps already benefit from Nextflow’s caching mechanism, implementing such differential processing for multi-trait steps would require custom state management beyond Nextflow’s intrinsic capabilities. Given this added complexity, these optimizations were not included in the current implementation but represent promising directions for future development.

Moreover, the current implementation supports the selection of only a single reference population across all traits. For large-scale studies, enabling trait-specific reference population selection would provide greater flexibility.

Finally, the exploration and visualization of stratified GWAS results, as supported in PheWeb 2 [7], are not yet available with GNExT. The ability to compare network medicine outcomes across stratified analyses would be of particular interest and represents a promising direction for future development.

In conclusion, GNExT represents a powerful tool for disseminating GWAS results. The nf-core pipeline streamlines data acquisition, normalization, updating, and analysis tasks underpinning the platform, scaling from a handful to thousands of traits. GNExT’s interactive web platform moves the field beyond the analysis of individual variants and bridges the gap to interpreting GWAS data through systems biology and network pharmacology.

## 4. Online Methods

### 4.1. Platform input

The primary input to GNExT’s preprocessing pipeline is a CSV phenotype file specifying the traits to be analyzed, including the file paths to the corresponding GWAS summary statistics. GWAS files based on either GRCh37 or GRCh38 genome assemblies can be used, since reference files are provided for both. The preprocessing pipeline additionally requires study-specific and computational configurations: the study-specific configurations define parameters such as the output directory, the structure of the GWAS summary statistics (via column indices), VEP annotation settings, and MAGMA parameters; the computational configurations specify memory and CPU allocations for each workflow component.

For the deployment of the GNExT platform after preprocessing, only a study-specific .env file must be defined. This file specifies absolute paths to the phenotype file, output directory of the preprocessing pipeline, the Typesense volume, and core service parameters such as the Typesense host and port. It also controls feature flags (e.g., activation of MAGMA components), theming parameters, study identifiers, and citation metadata. Optional predefined example traits, variants, and genes can be included to guide users during initial exploration. These configuration options ensure that the platform can be adapted to diverse research contexts while preserving a consistent and user-friendly interface. Full descriptions of all configuration parameters for both preprocessing and deployment are available in the corresponding GitHub repositories.

### 4.2. Harmonization of GWAS data formats

To accommodate diverse input formats, the first stage of the preprocessing pipeline harmonizes GWAS summary statistics. This step uses the Python package ZORP (https://github.com/statgen/zorp), which parses the GWAS data and abstracts format-specific differences. ZORP also exports the harmonized data as tab-separated, gzipped files, while variant sorting and indexing are handled in bash. The resulting harmonized dataset provides a standardized foundation for all downstream processing steps, ensuring independence from the original file structure.

### 4.3. Variant annotation using Ensembl VEP

Prior to variant annotation, the pipeline generates batch-specific VCF files for subsets of traits, which are subsequently merged into a single reference VCF containing all unique variants. This strategy remains efficient even when processing thousands of traits and ensures that each variant is annotated only once rather than repeatedly across individual GWAS files. For annotation, users may either supply a path to an existing local VEP cache or allow the pipeline to download it automatically. Cache retrieval and annotation rely on the nf-core modules ensemblvep download and ensemblvep vep [21]. We then populate two LMDB databases that function as on-demand lookup tables for the GNExT instances. The first LMDB database maps genetic variants to their corresponding rsIDs using the annotated VCF file. This avoids the need to append rsID annotations directly to every GWAS file, a process that can take several hours for large trait collections. A second LMDB database stores variant-to-gene mappings, recording for each variant the genes it overlaps or those located within a user-defined window. To ensure consistency, we use the gene coordinates of protein-coding genes from the same Ensembl release as VEP. For GRCh37 and GRCh38, gene annotation data is available for the two most recent Ensembl releases (114 and 115), and a script is provided to generate this data for other releases.

### 4.4. Features for result exploration

The generation of Manhattan and Q-Q plots within the GNExT framework builds upon the implementation provided by PheWeb, which produces the required visualization data in JSON format prior to platform deployment. Top genetic associations are extracted, whereas non-significant variants are aggregated into bins. This binning strategy substantially reduces the volume of data, compressing millions of variant-level values into manageable aggregates, thereby improving data transfer efficiency during API calls. The Top Hits table is generated by collecting all non-binned variants across traits, ranking them by *P*-value, and truncating the list according to a user-defined threshold. The resulting data are exported in JSON format for integration into the platform. Significance thresholds for Manhattan plots and criteria for defining Top Hits are fully configurable. Additionally, the pipeline produces BGZF-compressed and accompanying TBI-indexed files for each chromosome and each metric (*P*-value, allele frequency, effect size, and standard error). These indexed files support efficient variant-level data retrieval over all traits, required for PheWAS plots and variant-specific summary tables.

### 4.5. Gene-based association analysis

Gene-based association analysis is performed using MAGMA, aggregating single-variant summary statistics into gene-level *P*-values. We employ MAGMA’s default SNP-wise model, which uses the sum of squared SNP Z-statistics as the gene-level test statistic. To account for linkage disequilibrium (LD) among genetic variants, while avoiding the need for study-specific individual-level genotype data, LD structure is typically estimated using the 1000 Genomes Phase 3 reference dataset [1]. In this study, we generated harmonized and consistent reference data files for GRCh37 and GRCh38, which are deposited in a Zenodo repository.

MAGMA requires a gene annotation file specifying gene identifiers, chromosomal start and end coordinates, and strand information. Here, we use the same annotation source as employed for the LMDB variant-to-gene database. MAGMA subsequently performs an analogous internal positional mapping of variants to genes, assigning each variant to a gene when it falls within the gene boundaries or a predefined upstream or downstream window. Window parameters are held constant across the entire pipeline.

### 4.6. Disease module mining and drug repurposing with Drugst.One

GNExT integrates the Drugst.One interface, a plug-and-play web platform that converts systems biology methods into interactive tools for network analysis, disease mechanism mining, and drug repurposing. The interface consists of three components: a central network visualization window, a right-hand menu for direct network manipulation, and a left-hand panel system for node selection, task configuration and initiation, and result inspection. Analytical tasks operate on user-defined subnetworks, with results presented either directly in the Drugst.One interface or linking to external web pages. In GNExT, users can select a set of seed genes from the MAGMA results and transfer them into the integrated Drugst.One interface within the GNExT platform for the following downstream network-based analyses (for further details, we refer to the original Drugst.One and Drugst.One DREAM publications [42, 59]:

- Network integration: The selected seed proteins are mapped into a user-defined protein-protein interaction network, such as NeDRex, BioGRID, IID, IntAct, STRING, and APID, to investigate their interconnections. NeDRex is used by default, as it integrates all of these underlying databases into a unified interaction resource.
- Enrichment analysis: Gene set and pathway enrichment for any selected set of nodes can be performed using g:Profiler [55], while functional coherence assessment is available through DIGEST [2]. Drugst.One makes use of DIGEST’s reference-free mode, which quantifies internal functional coherence across all module genes, coherence is expressed as empirical *P*-values transformed on a −*log*_10_ scale, assuming that genes within a module are involved in similar biological processes. Both downstream analyses are accessible from the “Analysis” panel and automatically redirect to the corresponding web interfaces with the results preloaded.
- Disease module mining: The module mining functionality enables the detection of disease-associated protein modules within larger interactomes using dedicated disease module mining algorithms or centrality measures [42]. This analysis can be employed using the “Module detection” functionality within the “Analysis” panel. By default, a Multi-Steiner Tree approach is executed directly through the “Connect genes” panel.
- Drug prioritization: Seed and connector proteins can subsequently be evaluated as potential drug targets by querying established drug-protein interaction databases and ranking candidate compounds accordingly. This analysis is performed via the “Drug ranking” functionality in the “Analysis” panel, where users may select centrality-based metrics or TrustRank as the prioritization method.

As applied in both use cases, the “Quick drug search” option triggers the sequential execution of the Multi-Steiner tree module detection and harmonic centrality algorithm for drug prioritization [42].

### 4.7. Platform implementation

The preprocessing pipeline is implemented in Nextflow with support for local execution via Conda, containerized environments via Docker or Singularity, and distributed execution using SLURM-based HPC systems. The processed data are made accessible through a Django-based backend integrated with a Vue.js frontend. A Typesense search engine powers the platform’s autocomplete and query functionalities, enabling rapid and efficient retrieval of millions of genetic variants in real time. The entire system is containerized with Docker to ensure straightforward deployment, reproducibility, and portability across diverse computational environments.

### 4.8. Generation of reference genome data

MAGMA requires an external ancestry-appropriate reference panel in PLINK format (.bed, .fam, .bim) [53] to model LD among genetic variants when analyzing GWAS summary statistics. Many studies rely on the 1000 Genomes Project reference data [1], which is distributed in the appropriate format for genome assembly GRCh37 both within the MAGMA package and via FUMA. Since our workflow additionally supports inputs aligned to genome assembly GRCh38 and to ensure consistency across genome builds, we therefore generated all reference datasets uniformly for both GRCh37 and GRCh38.

Due to the immense file sizes of the GRCh38 build, all corresponding VCF files were first cleaned by restricting the “FORMAT” field to only genotypes, and by completely removing all entries in the “INFO” field. Following the FUMA preprocessing strategy, multi-allelic variants were decomposed into bi-allelic records using bcftools. For GRCh37, copy number variants were removed. For both genome builds, all chromosome-specific VCF files were then concatenated with bcftools. The resulting VCF was converted to PLINK binary format using PLINK2, assigning unique variant identifiers based on genomic coordinates (chr:pos:ref:alt) rather than rsIDs to ensure compatibility across different dbSNP [57] releases.

Finally, the resulting genotype data were stratified into five super-populations according to the provided panel file. The processed reference population files were deposited on Zenodo.

### 4.9. Application to GWAS olfaction data

To advance understanding of the genetic basis of human olfactory abilities, [23] performed a fixed-effect GWAS meta-analysis on odor identification using Sniffin’ Sticks across four cohorts totaling up to 21,495 individuals of European ancestry, including overall and sex-stratified models for 13 traits. A global identification score (range 0 to 12) summed correct identifications. The corresponding GWAS meta-analysis results (14 GB) were downloaded, preprocessed with the Nextflow pipeline. Gene definitions were obtained from Ensembl release 114 (GRCh38), and gene-level statistics were computed using the default 10kb up- and downstream window size in conjunction with the newly generated reference genome data aligned to GRCh38. For computational efficiency within the preprocessing pipeline, phenotypes were processed in batches of five.

### 4.10. Application to European-ancestry Pan-UK Biobank data

To demonstrate scalability of our GNExT ecosystem, a GNExT instance was deployed using the publicly available Pan-UK Biobank [31] GWAS summary statistics restricted to the European ancestry subset, comprising 7,160 traits.

The downloaded GWAS summary statistics (6.8 TB) were preprocessed to ensure a uniform file format and extract population-specific statistics, as not all input files adhered to the same structure due to case-control stratified reporting for binary phenotypes, which is incompatible with the GNExT ecosystem. Specifically, the allele frequencies for binary traits were derived by computing the weighted average of the corresponding case and control allele frequencies. Variants with allele frequencies equal to 0 or 1 were excluded, and *P*-values exceeding 99 and recorded as Infinity in the original input were capped at 99. Following preprocessing, the dataset was reduced to 2.1 TB. Gene annotations were obtained from Ensembl release 114 (GRCh37), and similar to the olfaction use case, gene-level statistics were computed using the default 10kb up- and downstream window size in combination with the newly generated GRCh37-aligned reference genome data. Phenotypes were processed in batches of 150.

## 5. Data and code availability

Summary statistics from the study by [23] were downloaded from https://www.health-atlas.de/assays/88 (accession code 8VK7H50F6P-6). Per-phenotype files for all available traits from the Pan-UK Biobank were downloaded from https://pan.ukbb.broadinstitute.org/downloads. Data of the 1,000 Genomes Project for GRCh37 and GRCh38 were downloaded from http://ftp.1000genomes.ebi.ac.uk/vol1/ftp/release/20130502/ and http://ftp.1000genomes.ebi.ac.uk/vol1/ftp/data_collections/1000G_2504_high_coverage/working/20201028_3202_raw_GT_with_annot/, respectively. The panel file specifying the samples super-populations was downloaded from https://ftp.1000genomes.ebi.ac.uk/vol1/ftp/release/20130502/integrated_call_samples_v3.20130502.ALL.panel. Gene annotation (GTF) files specifying gene coordinates were obtained from the corresponding Ensembl releases.

The source code of GNExT’s Nextflow-based preprocessing pipeline is available at https://github.com/DyHealthNet/gnext_nf_pipeline, the platform source code at https://github.com/DyHealthNet/gnext_platform. All reference files, including 1,000 Genomes PLINK files and the gene annotation files, are available on Zenodo (https://doi.org/10.5281/zenodo.17940903). The scripts to generate these reference files are available at https://github.com/DyHealthNet/gnext_reference_data. The GNExT instances are available at https://olfaction.gnext.gm.eurac.edu and https://panukbb-eur.gnext.gm.eurac.edu. The scripts and configuration files that were used to set up GNExT are available at the respective GitHub repositories.

## 6. Supplements

Supplementary data is available online.

## Author contributions statement

L.A. implemented the preprocessing pipeline, developed the platform together with support from B.R., and designed the Pan-UKBB use case. F.W. generated the reference datasets. L.A, F.W., and D.E. contributed to the olfaction use case, and J.F. assisted in interpreting the olfaction results. C.F., D.B.B., and M.L. obtained funding and supervised the project. All authors provided critical feedback and discussion, assisted in interpreting the results, writing the manuscript, and improving the web service.

## Acknowledgements

We thank Lisa Spindler and Andreas Maier for their continued support in integrating Drugst.One, and Daniele Di Domizio for his technical assistance with the deployment process.

