## Supplementary Material for "Bridging the gap between genome-wide association studies and network medicine with GNExT"

### 1 Supplementary Figures

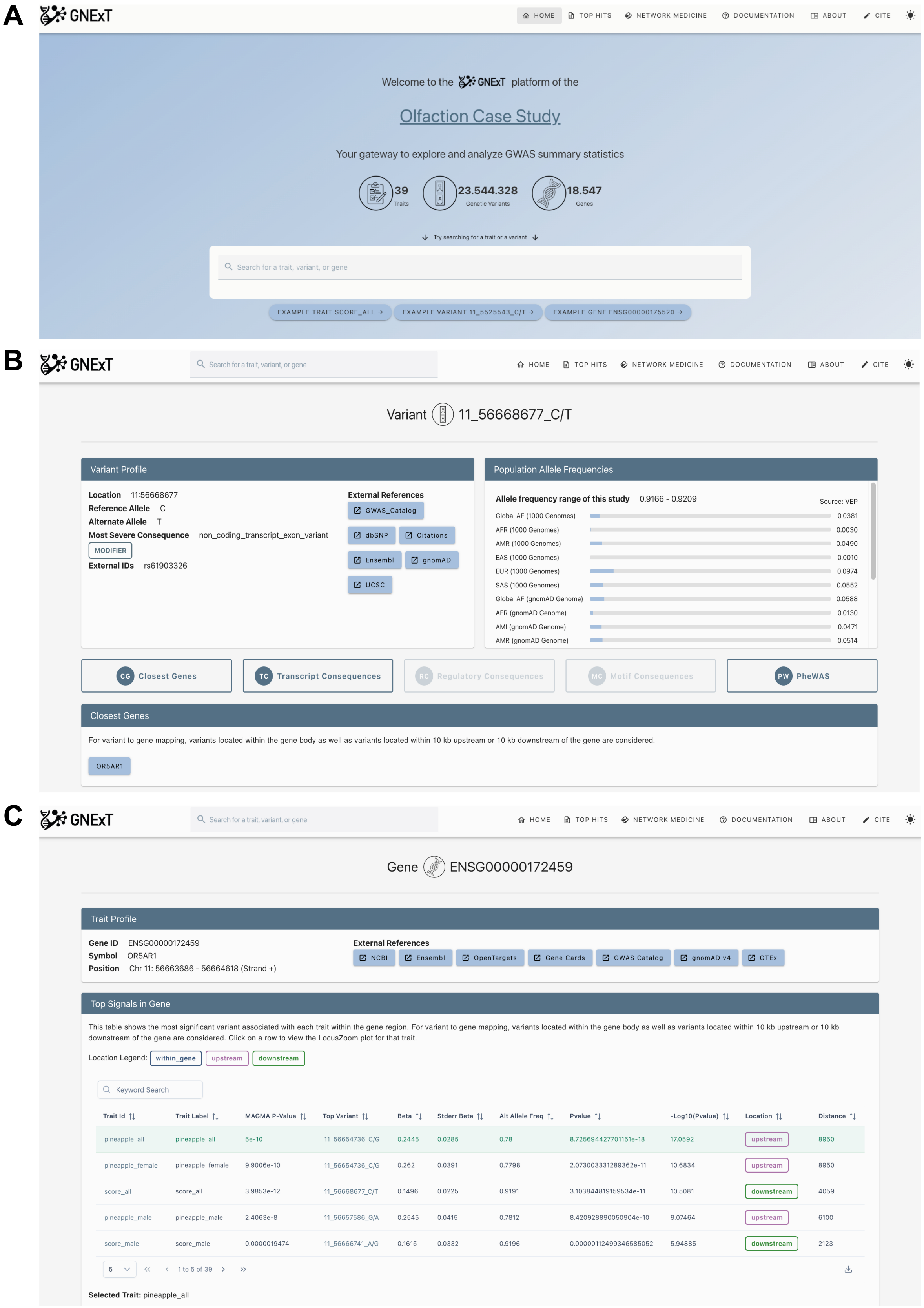

Supplementary Figure 1: Representative screenshots of the GNExT interface, showing the home page, variant-specific, and gene-specific page.

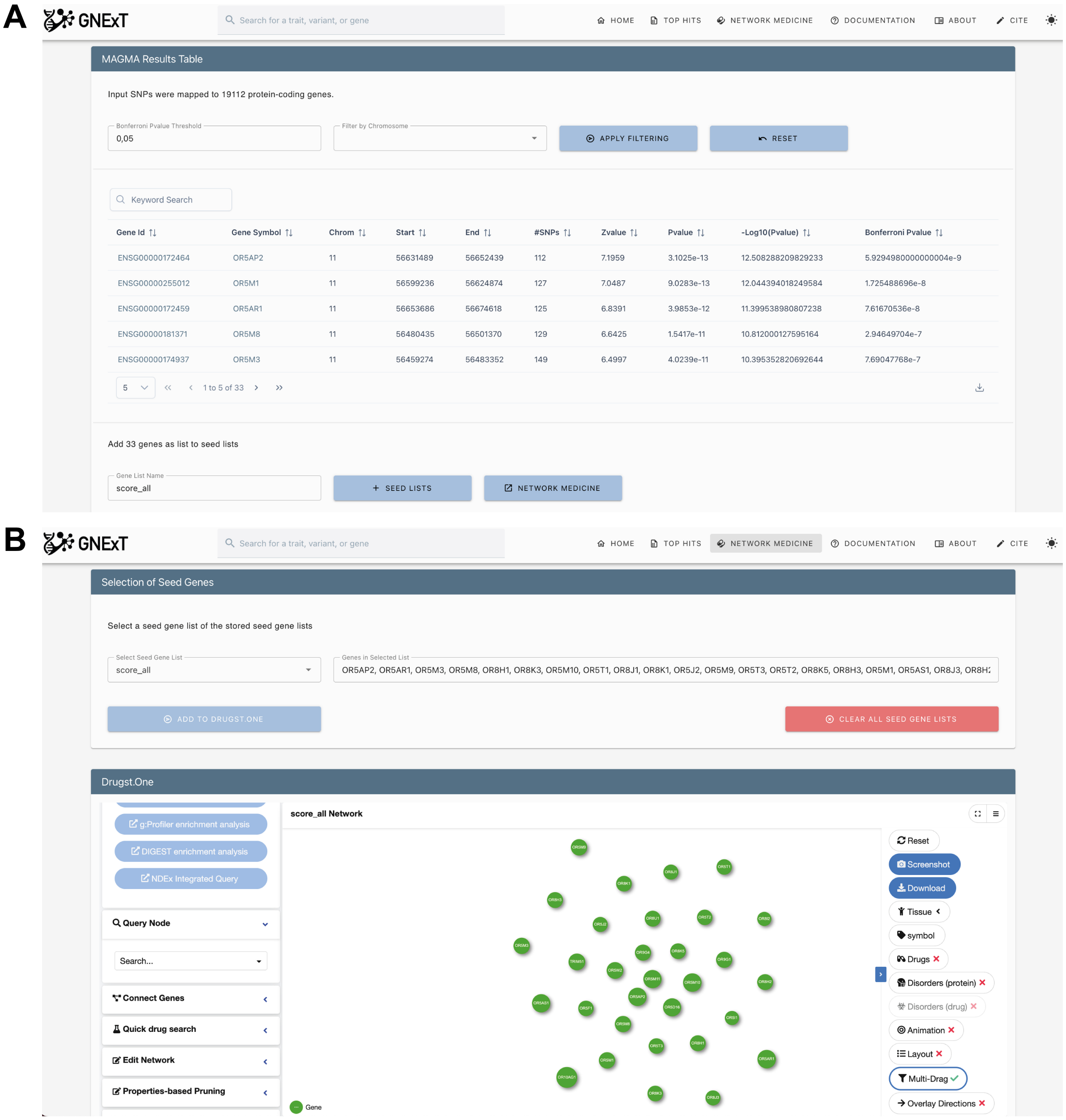

Supplementary Figure 2: Representative screenshots of the GNExT interface, highlighting the MAGMA results table on the trait-specific page and the network medicine page including the Drugst.One interface.

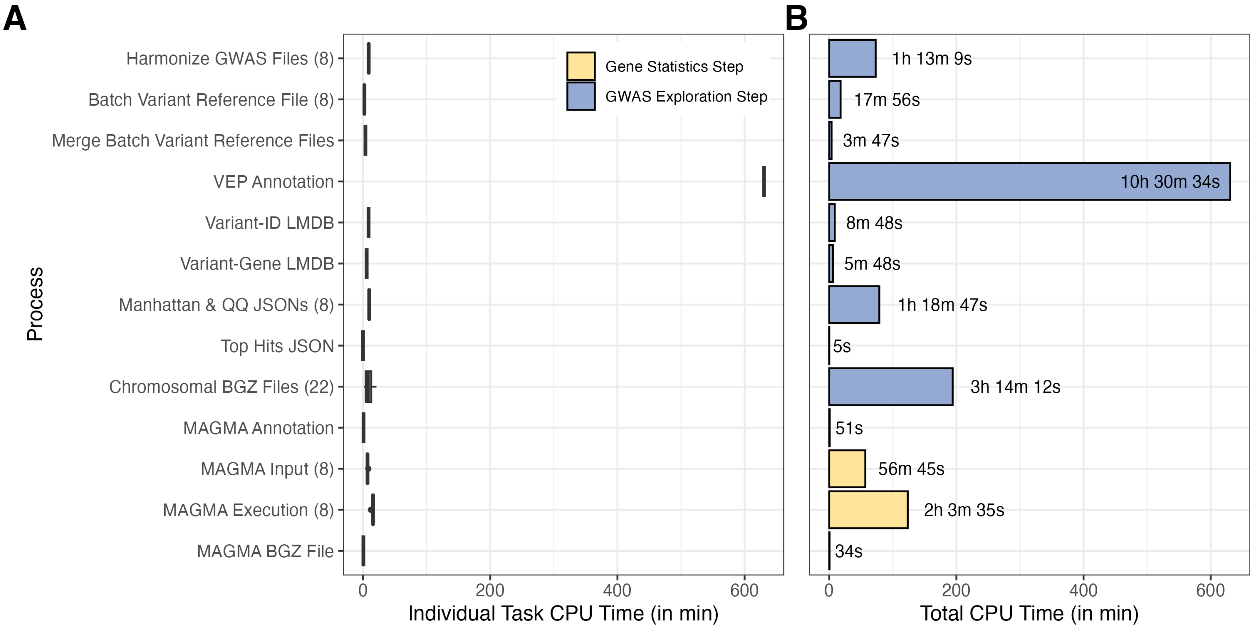

Supplementary Figure 3: Individual and total CPU time per process of the Nextflow preprocessing pipeline for the olfaction use case. Individual task CPU time was recorded intrinsically by Nextflow (A) and summed across all tasks belonging to each process (B). The x-axis denotes the process types, with the corresponding number of individual tasks per process arising from trait- or chromosome-batching shown in brackets.

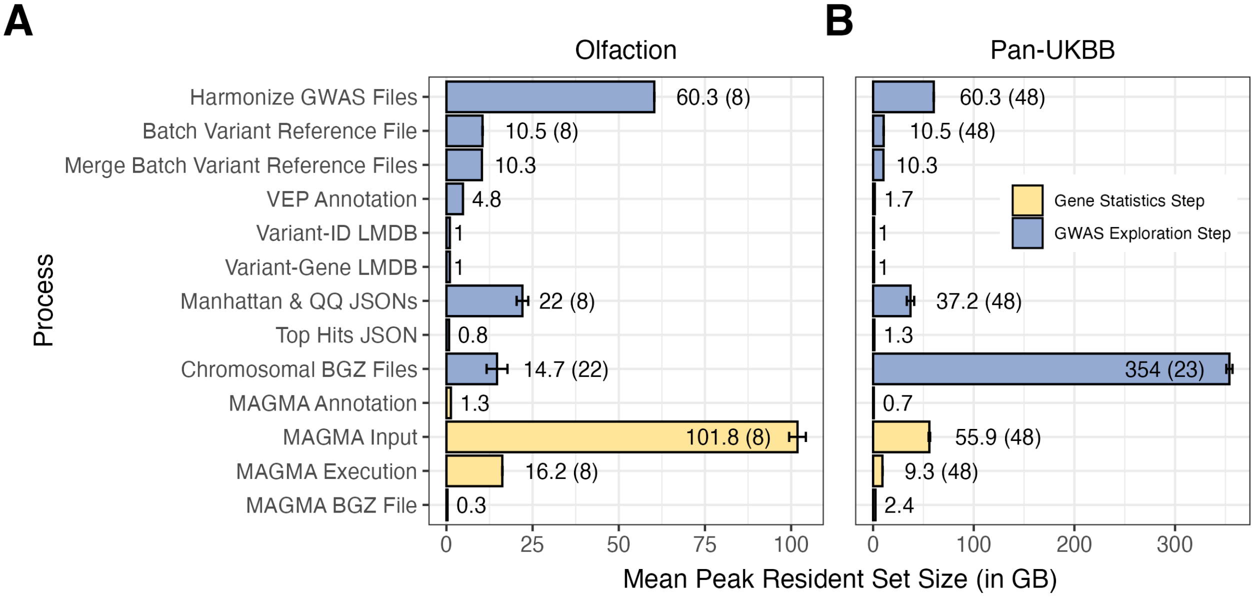

Supplementary Figure 4: Peak resident set size of the Nextflow preprocessing pipeline for the olfaction and Pan-UKBB use cases. Task-level peak resident set size was recorded natively by Nextflow for the olfaction (A) and Pan-UKBB use case (B), and results are summarized as mean ± standard deviation for each process type. The number of individual jobs per process arising from trait- or chromosome-batching are shown in brackets next to the memory label.

### 2 Supplementary Tables

Supplementary Table 1. MAGMA gene-based association results for the olfaction use case. 6

Supplementary Table 1: MAGMA gene-based association results for the olfaction use case. The "NeDReX" column indicates whether a given gene is present in the NeDReX database and can therefore be integrated into Drugst.One, whereas the "Module" column denotes whether the gene is part of the network module shown in Figure 3B.

| Symbol | ID | Chr | Start | End | #SNPs | Z-value | $P$-value | Bonferroni $P$-value | NeDReX | Module |
|---|---|---|---|---|---|---|---|---|---|---|
| OR5AP2 | ENSG00000172464 | 11 | 56631489 | 56652439 | 112 | 7.196 | $3.103 \times 10^{-13}$ | $5.929 \times 10^{-9}$ | yes | no |
| OR5M1 | ENSG00000255012 | 11 | 56599236 | 56624874 | 127 | 7.049 | $9.028 \times 10^{-13}$ | $1.725 \times 10^{-8}$ | yes | no |
| OR5AR1 | ENSG00000172459 | 11 | 56653686 | 56674618 | 125 | 6.839 | $3.985 \times 10^{-12}$ | $7.617 \times 10^{-8}$ | yes | yes |
| OR5M8 | ENSG00000181371 | 11 | 56480435 | 56501370 | 129 | 6.643 | $1.542 \times 10^{-11}$ | $2.946 \times 10^{-7}$ | yes | yes |
| OR5M3 | ENSG00000174937 | 11 | 56459274 | 56483352 | 149 | 6.500 | $4.024 \times 10^{-11}$ | $7.690 \times 10^{-7}$ | yes | yes |
| – | ENSG00000284732 | 11 | 56449221 | 56483352 | 204 | 6.355 | $1.045 \times 10^{-10}$ | $1.996 \times 10^{-6}$ | no | no |
| OR5T1 | ENSG00000181698 | 11 | 56264154 | 56286819 | 116 | 6.274 | $1.759 \times 10^{-10}$ | $3.362 \times 10^{-6}$ | yes | yes |
| OR8K3 | ENSG00000280314 | 11 | 56305144 | 56330639 | 131 | 6.244 | $2.127 \times 10^{-10}$ | $4.066 \times 10^{-6}$ | yes | no |
| OR8H1 | ENSG00000181693 | 11 | 56278462 | 56302254 | 123 | 6.214 | $2.580 \times 10^{-10}$ | $4.931 \times 10^{-6}$ | yes | no |
| OR5J2 | ENSG00000174957 | 11 | 56166618 | 56187556 | 110 | 6.181 | $3.183 \times 10^{-10}$ | $6.082 \times 10^{-6}$ | yes | no |
| OR8K5 | ENSG00000181752 | 11 | 56149394 | 56170317 | 103 | 6.169 | $3.444 \times 10^{-10}$ | $6.583 \times 10^{-6}$ | yes | no |
| OR5M11 | ENSG00000255223 | 11 | 56532340 | 56553257 | 133 | 6.153 | $3.794 \times 10^{-10}$ | $7.251 \times 10^{-6}$ | yes | yes |
| OR5M10 | ENSG00000254834 | 11 | 56566774 | 56587721 | 122 | 6.150 | $3.884 \times 10^{-10}$ | $7.424 \times 10^{-6}$ | yes | yes |
| OR5M9 | ENSG00000150269 | 11 | 56452469 | 56473401 | 119 | 6.109 | $5.000 \times 10^{-10}$ | $9.556 \times 10^{-6}$ | yes | no |
| OR8J1 | ENSG00000172487 | 11 | 56344291 | 56371511 | 155 | 6.109 | $5.000 \times 10^{-10}$ | $9.556 \times 10^{-6}$ | yes | no |
| OR8J2 | ENSG00000254658 | 11 | 56198985 | 56225222 | 141 | 6.109 | $5.000 \times 10^{-10}$ | $9.556 \times 10^{-6}$ | no | no |
| OR5T3 | ENSG00000172489 | 11 | 56242254 | 56263222 | 113 | 6.109 | $5.000 \times 10^{-10}$ | $9.556 \times 10^{-6}$ | yes | no |
| OR8K1 | ENSG00000150261 | 11 | 56336039 | 56356998 | 102 | 6.109 | $5.000 \times 10^{-10}$ | $9.556 \times 10^{-6}$ | yes | no |
| OR5T2 | ENSG00000181718 | 11 | 56221282 | 56244255 | 108 | 6.060 | $6.817 \times 10^{-10}$ | $1.303 \times 10^{-5}$ | yes | no |
| OR8H3 | ENSG00000181761 | 11 | 56112373 | 56133311 | 185 | 5.976 | $1.144 \times 10^{-9}$ | $2.187 \times 10^{-5}$ | yes | no |
| OR8J3 | ENSG00000167822 | 11 | 56124721 | 56150201 | 163 | 5.938 | $1.442 \times 10^{-9}$ | $2.757 \times 10^{-5}$ | yes | no |
| OR10AG1 | ENSG00000174970 | 11 | 55955755 | 55979945 | 261 | 5.823 | $2.895 \times 10^{-9}$ | $5.533 \times 10^{-5}$ | yes | yes |
| OR5AS1 | ENSG00000181785 | 11 | 56017654 | 56048191 | 376 | 5.797 | $3.379 \times 10^{-6}$ | $6.458 \times 10^{-5}$ | yes | yes |
| OR8H2 | ENSG00000181767 | 11 | 56093687 | 56117658 | 257 | 5.787 | $3.586 \times 10^{-9}$ | $6.853 \times 10^{-5}$ | yes | yes |
| OR8I2 | ENSG00000172154 | 11 | 56083308 | 56104240 | 220 | 5.715 | $5.482 \times 10^{-9}$ | $1.048 \times 10^{-4}$ | yes | no |
| OR5F1 | ENSG00000149133 | 11 | 55983681 | 56004625 | 236 | 5.634 | $8.788 \times 10^{-9}$ | $1.680 \times 10^{-4}$ | yes | yes |
| OR5W2 | ENSG00000187612 | 11 | 55903650 | 55924582 | 212 | 5.559 | $1.358 \times 10^{-5}$ | $2.595 \times 10^{-4}$ | yes | no |
| OR5I1 | ENSG00000167825 | 11 | 55925456 | 55946400 | 178 | 5.542 | $1.497 \times 10^{-8}$ | $2.862 \times 10^{-4}$ | yes | no |
| OR9G4 | ENSG00000172457 | 11 | 56731223 | 56758697 | 169 | 5.302 | $5.743 \times 10^{-8}$ | $1.098 \times 10^{-3}$ | yes | no |
| OR5G3 | ENSG00000241356 | 11 | 56809573 | 56830516 | 145 | 5.256 | $7.370 \times 10^{-8}$ | $1.409 \times 10^{-3}$ | no | no |
| TRIM51 | ENSG00000124900 | 11 | 55873297 | 55901810 | 210 | 5.252 | $7.506 \times 10^{-8}$ | $1.434 \times 10^{-3}$ | yes | yes |
| OR9G1 | ENSG00000174914 | 11 | 56689095 | 56713884 | 142 | 4.964 | $3.448 \times 10^{-7}$ | $6.589 \times 10^{-3}$ | yes | yes |
| OR8U1 | ENSG00000172199 | 11 | 56365624 | 56386553 | 129 | 4.913 | $4.486 \times 10^{-4}$ | $8.574 \times 10^{-3}$ | yes | yes |

Supplementary Table 2: Drug prioritization results for the olfaction network module presented in Figure 3B.

| Label | ID | DrugBank ID | Score | Rank | Status |
|---|---|---|---|---|---|
| Sunitinib | dr1247 | DB01268 | 1.000 | 1 | approved |
| Fostamatinib | dr10037 | DB12010 | 1.000 | 1 | approved |
| Nintedanib | dr8092 | DB09079 | 1.000 | 1 | approved |
| Midostaurin | dr5745 | DB06595 | 1.000 | 1 | approved |
| Fedratinib | dr10511 | DB12500 | 1.000 | 1 | approved |
| Crizotinib | dr7891 | DB08865 | 1.000 | 1 | approved |
| Bosutinib | dr5762 | DB06616 | 9.169 | 2 | approved |
| Vandetanib | dr4939 | DB05294 | 9.169 | 2 | approved |
| Pacritinib | dr9732 | DB11697 | 8.982 | 3 | approved |
| Pexidartinib | dr10967 | DB12978 | 8.982 | 3 | approved |
| Entrectinib | dr10013 | DB11986 | 8.982 | 3 | approved |
| Tivozanib | dr9832 | DB11800 | 8.982 | 3 | approved |
| Erlotinib | dr516 | DB00530 | 8.982 | 3 | approved |
| Sorafenib | dr386 | DB00398 | 8.982 | 3 | approved |
| Axitinib | dr5770 | DB06626 | 8.982 | 3 | approved |
| Dasatinib | dr1234 | DB01254 | 8.982 | 3 | approved |
| Tofacitinib | dr7920 | DB08895 | 8.305 | 4 | approved |
| Lenvatinib | dr8091 | DB09078 | 8.305 | 4 | approved |
| Selpercatinib | dr13602 | DB15685 | 8.305 | 4 | approved |
| Palbociclib | dr8086 | DB09073 | 8.004 | 5 | approved |
| Regorafenib | dr7921 | DB08896 | 8.004 | 5 | approved |
| Ponatinib | dr7926 | DB08901 | 8.004 | 5 | approved |
| Afatinib | dr7940 | DB08916 | 8.004 | 5 | approved |
| Imatinib | dr605 | DB00619 | 8.004 | 5 | approved |
| Pralsetinib | dr13736 | DB15822 | 8.004 | 5 | approved |
| Quizartinib | dr10867 | DB12874 | 8.004 | 5 | approved |
| Pazopanib | dr5741 | DB06589 | 8.004 | 5 | approved |
| Brigatinib | dr10287 | DB12267 | 8.004 | 5 | approved |
| Neratinib | dr9859 | DB11828 | 8.004 | 5 | approved |
| Ibrutinib | dr8067 | DB09053 | 8.004 | 5 | approved |
| Infigratinib | dr9917 | DB11886 | 8.004 | 5 | approved |
| Gilteritinib | dr10166 | DB12141 | 8.004 | 5 | approved |
| Trilaciclib | dr13364 | DB15442 | 8.004 | 5 | approved |
| Filgotinib | dr12785 | DB14845 | 8.004 | 5 | approved |
| Ceritinib | dr8076 | DB09063 | 8.004 | 5 | approved |
| Nilotinib | dr4584 | DB04868 | 8.004 | 5 | approved |
| Gefitinib | dr307 | DB00317 | 8.004 | 5 | approved |
| Cabozantinib | dr7900 | DB08875 | 8.004 | 5 | approved |
| Acetic acid | dr2981 | DB03166 | 7.462 | 6 | approved |

Supplementary Table 3: MAGMA gene-based association results for the Pan-UKBB Alzheimer's disease use case. The "NeDReX" column indicates whether a given gene is present in the NeDReX database and can therefore be integrated into Drugst.One, whereas the "Module" column denotes whether the gene is part of the network module shown in Figure 5G. PVRL2 was manually added to the Drugst.One interface as NECTIN2 using the "Edit Network" functionality.

| Symbol | ID | Chr | Start | End | #SNPs | Z-value | $P$-value | Bonferroni $P$-value | NeDReX | Module |
|---|---|---|---|---|---|---|---|---|---|---|
| TOMM40 | ENSG00000130204 | 19 | 45383826 | 45416946 | 152 | 7.662 | $9.159 \times 10^{-15}$ | $1.837 \times 10^{-10}$ | yes | yes |
| PVRL2 | ENSG00000130202 | 19 | 45339432 | 45402485 | 384 | 7.123 | $5.281 \times 10^{-13}$ | $1.059 \times 10^{-8}$ | yes | yes |
| APOE | ENSG00000130203 | 19 | 45399011 | 45422650 | 98 | 6.109 | $5.000 \times 10^{-10}$ | $1.003 \times 10^{-5}$ | yes | yes |
| APOC1 | ENSG00000130208 | 19 | 45407504 | 45432606 | 95 | 6.109 | $5.000 \times 10^{-10}$ | $1.003 \times 10^{-5}$ | yes | yes |
| BCAM | ENSG00000187244 | 19 | 45302328 | 45334673 | 168 | 5.128 | $1.462 \times 10^{-7}$ | $2.934 \times 10^{-3}$ | yes | yes |
| RELB | ENSG00000104856 | 19 | 45494688 | 45551452 | 205 | 4.882 | $5.248 \times 10^{-7}$ | $1.053 \times 10^{-2}$ | yes | yes |
| AC005779.2 | ENSG00000267545 | 19 | 45673080 | 45715702 | 242 | 4.860 | $5.858 \times 10^{-7}$ | $1.175 \times 10^{-2}$ | no | no |
| CLPTM1 | ENSG00000104853 | 19 | 45447842 | 45506599 | 260 | 4.795 | $8.135 \times 10^{-7}$ | $1.632 \times 10^{-2}$ | yes | yes |
| MARK4 | ENSG00000007047 | 19 | 45572546 | 45818541 | 1137 | 4.718 | $1.193 \times 10^{-6}$ | $2.393 \times 10^{-2}$ | yes | yes |
| AC006126.3 | ENSG00000266958 | 19 | 45680646 | 45730191 | 316 | 4.677 | $1.453 \times 10^{-6}$ | $2.915 \times 10^{-2}$ | no | no |
| EXOC3L2 | ENSG00000130201 | 19 | 45705879 | 45747469 | 265 | 4.612 | $1.997 \times 10^{-6}$ | $4.006 \times 10^{-2}$ | yes | no |
| NVL | ENSG00000143748 | 1 | 224405036 | 224528089 | 392 | 4.576 | $2.365 \times 10^{-6}$ | $4.744 \times 10^{-2}$ | yes | yes |

Supplementary Table 4: Drug prioritization results for the Alzheimer's network module presented in Figure 5G.

| Label | ID | DrugBank ID | Score | Rank | Status |
|---|---|---|---|---|---|
| Metformin | dr320 | DB00331 | 1.000 | 1 | approved |
| Chlorzoxazone | dr345 | DB00356 | 1.000 | 1 | approved |
| Sunitinib | dr1247 | DB01268 | 9.698 | 2 | approved |
| Fostamatinib | dr10037 | DB12010 | 9.698 | 2 | approved |
| Nintedanib | dr8092 | DB09079 | 9.698 | 2 | approved |
| Midostaurin | dr5745 | DB06595 | 9.698 | 2 | approved |
| Fedratinib | dr10511 | DB12500 | 9.698 | 2 | approved |
